# Optimized immunoglobulin knock-ins using Cas9 reveal peritoneal B cell lineage relationships *in vivo*

**DOI:** 10.1101/2021.10.31.466662

**Authors:** Steven A. Erickson, Eliot Zapata-Bultman, Linda Degenstein, Albert Bendelac

**Author notes:** **Correspondence to:** Albert Bendelac.

## Abstract

Immunoglobulin (Ig) knock-in mice are valuable tools in basic and translational immunological research. Here we present “Speed-Ig,” a rapid Cas9-based method for generating Ig knock-in mouse lines with high on-target integration rates at both heavy and light chain alleles. With standardized target sites and promoter regions, Speed-Ig mice can be used for comparative studies of B cell biology and vaccine optimization *in vivo*. We used Speed-Ig to create panels of mice with Ig pairs derived from B-1a, B-1b, and B-2 cells. Surprisingly, B-1b and B-2 Ig pairs drove both B-1b and B-2 phenotypes, suggesting a previously unknown lineage relationship between these subsets. We then confirmed the B-1:B-2 relationship with transcription factor reporter lines and through adoptive cell transfer experiments. In summary, our Ig knock-in approach facilitated the discovery of previously unappreciated aspect of innate-like B cell biology.

## INTRODUCTION

Mammalian humoral immune responses are critical in host protection against pathogen challenge. B lymphocytes, the chief effector cells of humoral immunity, produce antibodies which facilitate recognition and clearance of pathogens in an antigen-specific manner. Eliciting protective humoral responses toward currently intractable infections, such as human immunodeficiency virus and influenza virus, is a major goal of current vaccine research.^1–4^ In addition to combating infections, antibodies are being used therapeutically to treat a variety of human diseases.^5^ Moreover, B cells and their antibody autoimmune pathologies, allergic responses, and organismal homeostasis.^6,7^ Understanding the development and differentiation of B cells bearing defined immunoglobulin (Ig) pairs therefore holds tremendous value in basic and translational biological research. However, underlying population diversity in wild-type animals confounds analysis of clonal B cell behavior *in vivo*. The pioneering work of Klaus Rajewsky and colleagues developed targeted Ig integrations in mice such that virtually all B cells solely express the inserted Ig chains.^8^ Monoclonal Ig knock-ins preserve Ig locus-dependent features including affinity maturation and class switch recombination and thus closely approximate natural B cell biology, but are costly and time-consuming to create.^9^ These constraints may limit the scope of knock-in studies, and potentially lead to underrepresentation of heterogeneity in the B cell subsets being analyzed. Modern gene editing techniques incorporating programmable nucleases can greatly accelerate the generation of fixed-Ig mice while reducing costs, expediting the *in vivo* analysis of large Ig knock-in line panels.^10,11^ This is of direct relevance to understudied B cell populations such as those found in the peritoneal and pleural cavities of mice.

Innate-like B cells, in particular the B-1 lineages in mice, are thought to provide rapid, non-specific defense against infection and constitutive protection from autoimmunity^12–14^. Primarily located in the cavities surrounding the lungs and intestine, B-1 cells phenotypically differ from conventional B cells in several ways. Both CD5^+^ B-1a and CD5^-^ B-1b cells are larger and more granular than B-2 cells and have a distinct set of surface markers^15^. Several transcription factors, including *Basic helix loop helix family, member e41* (*Bhlhe41)* and *Zinc finger and BTB domain-containing protein 32* (*Zbtb32)*, have been associated with the B-1 phenotype^16–18^.The majority of B-1 studies have focused on B-1a cells, which are at least in part seeded by precursors in early life and show a strong repertoire bias^19–22^. The temporal dynamics of B-1a ontogeny evoked two competing but not necessarily exclusive models of B-1 cell differentiation^23^. Succinctly, the “lineage” hypothesis predicts independent precursors for B-1 and B-2 cells, while the “induced differentiation” hypothesis predicts Ig-dependent cell fating from otherwise dually potent precursors. Transgenic and knock-in mice bearing “canonical” V_H_11/V_κ_9 or V_H_12/V_κ_4 Ig pairs drive strong B-1a lineage biases^24,25^. Though clearly an Ig-dependent process, these Ig pairs typically recombine in fetal/neonatal progenitors. Modulation of BCR signaling strength alone can alter subset segregation^26–28^. Additionally, analysis of cells from a Nur77 reporter line, serving as a proxy for BCR signaling, indicate that B-1a (but not B-1b) have higher steady state BCR stimulation than B-2 cells^29^. Recently, a mouse model in which mature B-2 cells are induced to switch expression from a B-2 Ig pair to one used by canonical B-1a cells, differentiation to the B-1a lineage was observed^18^. However, studies on canonical B-1a Ig pairs only account for a fraction of the B-1a repertoire and none of the B-1b subset, of which very little is understood besides a lack of surface CD5 expression. Whether non-canonical B-1a or B-1b cell biology support the lineage or induced differentiation hypotheses is an open question.

Here, we present “Speed-Ig,” a rapid and efficient method for generating Ig knock-in mice. Through microinjection of single-cell murine embryos with donor DNA, gRNA, and Cas9 protein followed by a single breeding, paired-chain monoclonal Ig mouse lines were developed in less than three months. Ig knock-in mice displayed robust Ig expression, B cell development, and subset differentiation. Our method serves as a platform for testing the development and differentiation of B cells directed by Ig pairs of interest. Using Speed-Ig, we observed that non-canonical B-1a Ig pairs, much like canonical B-1a receptors, imprint a B-1a phenotype in expressing cells. However, we observed unexpected phenotypic flexibility in B-1b and B-2 Ig knock-in mice. Both B-1b and B-2 cells were observed in either panel of knock-in lines, suggesting that factors outside of BCR signaling may be involved in fate decisions. Strikingly, a large fraction of peritoneal B cells in B-1b and B-2 knock-in lines took on a phenotype intermediate to either subset, potentially representing a transitional stage. Using a *Zbtb32*^*EGFP*^ reporter line, we observed that B-1/B-2 polarity correlates with Zbtb32 expression. Furthermore, in a fate-mapping approach using *Bhlhe41*^*Cre*^, a third of peritoneal B-2 cells and nearly all B-1/B-2 intermediate cells were labeled. Adoptive transfers revealed that both B-2 and intermediate cells can become bona fide B-1 cells, with little plasticity from B-1a/B-1b to B-2. Thus, we report additional complexity to B-1 cell differentiation not fully explained by existing models and propose that additional layers of B-1 subset seeding exist.

## MATERIALS AND METHODS

### Mice

C57BL/6J, B6.SJL-*Ptprc*^*a*^*Pepc*^*b*^/BoyJ, *CD23*^*Cre*^ (B6.Cg-Tg(Fcer2a-cre)5Mbu/J)^30^, *hCk* (B6;129-Igkctm1(IGKC)Mnz/J)^31^, *IgHa* (B6.Cg-Gpi1a Thy1a Igha/J), *R26R-EYFP* (B6.129×1-Gt(ROSA)26Sortm1(EYFP)Cos/J)^32^ and *Rag1*^*-/-*^ (B6.129S7-Rag1tm1Mom/J) mice were purchased from The Jackson Laboratory. *Rag2-EGFP* (FVB-Tg(Rag2-EGFP)1Mnz/J)^33^ were purchased from the Jackson Laboratory and backcrossed to C57BL/6J. Ig knock-ins, the *Zbtb32*^*EGFP*^ reporter, and the *Bhlhe41*^*Cre*^ lines were generated for this study. Mice deficient for the IgH J segment loci were generated for this study using Cas9 and the following guide RNA sites: GCTACTGGTACTTCGATGTC and GCCATTCTTACCTGAGGAGA. For experiments, mice were analyzed at 3-8 weeks and compared to littermate controls. Mice were maintained in a specific pathogen-free environment at the University of Chicago. All experimental guidelines were approved by the Institutional Animal Care and Use Committee (IACUC).

### Single-cell cloning of rearranged Ig genes

Single B cells were sorted as previously described^34–36^ from the following populations using the indicated markers: Splenic B-2 (B220^+^ CD19^+^ CD21^+^ CD23^+^), peritoneal B-1a (CD19^+^ CD23^-^ CD5^+^), and peritoneal B-1b (CD19^+^ CD23^-^ CD5^-^). Briefly, single cells were sorted into 96-well plates (Biorad # HSP9601) containing 10μl of 1% v/v β-mercaptoethanol TCL buffer (Qiagen # 1031576) using a BD AriaII cell sorter. RNA was purified using RNAse-free SPRI beads^37,38^ (GE Healthcare #65152105050250) and followed by cDNA synthesis using Superscript IV cDNA synthesis kit (Invitrogen #18091050) as described^36^. Rearranged Ig chains were amplified and sequenced using degenerate Ig primers^35^. In-frame variable regions were screened in IMGT^39^ then amplified with gene-specific overlap primers for subsequent Gibson assemblies^40^ (**Supplementary Table 1**).

### Guide site selection

Guide sites were identified using Integrated DNA Technologies’ (IDT) selection tool (https://www.idtdna.com/site/order/designtool/index/CRISPR_CUSTOM) by inputting ∼500bp surrounding the region of interest. Individual guide sites were selected by a number of criteria, including on-target scores, off-target scores and proximity to the terminal stop codon (for non-Ig knock-ins). In general, on-target scores held primacy in selection, as off-target scores are skewed by non-PAM-led genomic hits and off-target integrations can be easily bred out unless falling near the intended target. Similar criteria were used when evaluating guide site options from CHOPCHOP (https://chopchop.cbu.uib.no/) and the Broad sgRNA designer (https://portals.broadinstitute.org/gpp/public/analysis-tools/sgrna-design).

### Donor DNA construct generation

For Speed-Ig Donor DNA constructs, the pUC19 plasmid was digested with SacI and SbfI. Linearized plasmids were then assembled with overlapping homology arms encompassing both 5’ and 3’ arms to generate ‘intermediate’ targeting constructs. Promoter regions were amplified from genomic DNA (**Supplementary Table 1**), and full targeting constructs were obtained by assembling SalI-linearized intermediate constructs, promoter regions, and Ig variable regions. All assemblies were performed with NEB HiFi 2X Master Mix (NEB #E2621L). Assembled constructs were transformed into XL-10 Gold Ultracompetent *E. coli* (Agilent #200314). Colonies were miniprepped and Sanger sequenced using a set of primers to ensure proper assembly and in-frame V regions (**Supplementary Table 1**). Glycerol stocks of desired clones were midiprepped using anion-exchange columns (Invitrogen #K210015) following growth in 50ml LB + carbenicillin medium for 10-12 hours. Midiprepped DNA was then ethanol precipitated, resuspended in nuclease-free water, and passed through 0.1um filters (EMD Millipore #UFC30VV25). DNA concentration was measured by spectrophotometry on a NanoDrop™; preparations falling below 1.8 A_260_/A_280_ were discarded.

### Injection mix preparation

Three hours prior to microinjection, tracrRNA (IDT #1072532) and crRNA (IDT) were annealed and complexed with Cas9 protein (IDT #1081058) as described^41^. Briefly, 5μl of 1μg/μl cRNA was incubated with 10μl of 1μg/μl tracrRNA at 95°C for five minutes followed by cooling to 25°C with ramp speed of 5°C min^-1^. In a sterile nuclease-free tube (Eppendorf #022600001), 5μl of annealed crRNA:tracrRNA was then complexed with 5μl of 1μg/μl Cas9 protein at 25°C for 15 minutes in 50μl nuclease-free ultrapure water (Thermo #J71786XCR). Following gRNA:Cas9 complexing, donor DNA was added along with ultrapure water to give a final volume of 100μl and concentrations of 50ng/μl crRNA:tracrRNA (or sgRNA), 50ng/μl Cas9 protein (or 125ng/μl mRNA), and 15ng/μl targeting construct DNA. The complete injection mix was spun at 20,000 x *g* for ten minutes, and the top 65μl was transferred to a new sterile tube for transport to Transgenics core facilities. For injections with sgRNA and Cas9 mRNA, respective plasmids were linearized and transcribed in vitro prior to injection as described^42^.

### Embryo microinjection and transgenesis

C57BL/6J female mice in diestrus between the age of 9 and 16 weeks of age were selected for superovulation. Depending on age and size of the mice, 5-7 IU of PMSG (Millipore #367222/BioVendor #RP1782721000) was injected (IP) at 2:50-3:05PM, followed by HCG 5-7 IU (Sigma CG5) 47 hrs. later. Mice were paired with C57BL/6J males one hour later. Females were checked the next morning for evidence of a mating plug; plugged mice were harvested for fertilized eggs. Eggs were cultured for 1-3 hrs. in KSOMaa Evolve Bicarb +BSA (Zenith Biotech #ZEKS-050). Glass needles were loaded with the described injection mix and placed on ice until used for injection. Eggs were placed on the injection slide in KSOMaa Evolve HEPES +BSA (Zenith Biotech #ZEHP-050). Fertilized eggs were injected using an Eppendorf FemtoJet apparatus. Injected eggs were returned to the incubator and cultured until embryo transfer into 0.5d.e. pseudopregnant CD-1 females. Implanted females were observed daily and separated 2 days before the pups were due to deliver. Tail samples were taken between 10 and 15 days for genotyping.

### Ig knock-in genotyping

Tail clippings from potential founder animals were taken for genotyping. DNA was extracted in 300μl 50mM NaOH for 30’ at 95°C. Samples were neutralized with 25μl 1M Tris-HCl pH7.5 and used as template DNA for PCR with targeting-specific primers (**Supplementary Table 1**). Reactions were carried out with OneTaq 2X master mix (NEB #M0488S) according to manufacturer instructions.

### Southern blotting

DNA was extracted from spleen segments using DNeasy Blood & Tissue Kit (Qiagen #69504). 15ug of DNA were digested overnight with the indicated restriction enzymes and run on 0.8% agarose gel. Gels were denatured in 0.5M NaOH 1.5M NaCl with gentle agitation for 15 minutes twice before neutralization in 0.5M Tris-HCl pH 7.5 1.5M NaCl for 15 minutes twice. DNA was transferred from gels to positively charged nylon membranes (Invitrogen #AM10102) overnight via capillary transfer in 20X SSC (Thermo # 15557036). Prior to transfer, nylon membranes were soaked in nanopure water for five minutes followed by 20X SSC for five minutes. Following transfer, nylon membranes were soaked in 2X SSC for five minutes followed by UV cross-linking. Cross-linked membranes were blocked in pre-warmed DIG EasyHyb (Roche # 11603558001) at 43°C for four hours rolling in hybridization tubes. Custom DIG-labeled probes were generated using PCR DIG Probe Synthesis Kit (Roche #11636090910) and blots were hybridized for 16 hours at 48°C with a final concentration of 25ng/ml probe in 10ml DIG EasyHyb buffer rolling in hybridization tubes. Following hybridization, membranes were washed twice with low stringency buffer (0.5X SSC 0.1% SDS) at 25°C for five minutes, then twice with high stringency buffer (2X SSC 0.1% SDS) at 68°C for fifteen minutes. Membranes were equilibrated in wash buffer (Roche #11585762001) at 25°C for five minutes with gentle shaking, then blocked in 100ml block buffer (Roche #11585762001) at 25°C for two hours with gentle shaking. Hybridized probe was detected using AP-conjugated anti-DIG Fab fragments (Roche #11093274910) via incubation at 1:10,000 in 50ml at 25°C for thirty minutes with gentle shaking. Membranes were washed three times in wash buffer at 25°C for fifteen minutes. Membranes were equilibrated (Roche #11093274910) and CDP-Star Substrate (Invitrogen #T2145) was used to detect AP.

### Lymphocyte isolation

Spleen and bone marrow were collected and passed through 70 μm cell strainers, centrifuged, and resuspended in flow cytometry buffer. For peritoneal lavage, 5ml PBS 3% FBS was injected with a 30g needle into the peritoneal cavity followed by gentle shaking. Fluid was then collected with a 25g needle, centrifuged, and resuspended in flow cytometry buffer. Small intestinal lamina propria isolation was performed as follows: tissues were excised and fat, Peyer’s patches, and intestinal contents removed. Tissues were opened longitudinally, cut into ∼1 cm pieces, and placed into a 50 mL tube with 10 mL RPMI 1% FCS 1 mM EDTA. Small intestines were processed as two halves in separate tubes and combined after lamina propria lymphocyte isolation as noted below. Samples were incubated at 37°C with shaking for 15 min. Pieces were collected on a 100 μm cell strainer (Fisher) and placed in fresh RPMI 1% FCS 1mM EDTA for another 15 min with shaking at 37°C. Pieces were again collected on a 100 μm cell strainer and flow-through was discarded. Intestinal pieces were then placed into 10 mL RPMI 20% FCS with 0.5 mg/mL Collagenase Type I (Gibco #17100017) and 0.1 mg/mL DNase I (Sigma #DN25) at 37°C with shaking for 30 min. Supernatant was collected by filtering through a fresh 100 μm cell strainer and intestinal pieces were placed in fresh RPMI 20% FCS with collagenase and DNase for another 30 min with shaking at 37°C. Supernatant was again collected, remaining tissues mashed, and the cell strainer was washed with 30 mL ice-cold RPMI. The two lamina propria lymphocyte fractions were combined after centrifugation and further enriched by centrifugation in 40% Percoll^®^ (Sigma #P1644) to remove epithelial cells and debris, washed in flow cytometry buffer, and resuspended for cell staining.

### Flow cytometry

Cells were incubated with anti-CD16/32 (Fc block) prior to staining with the following fluorophore or biotin conjugated monoclonal antibodies purchased from Biolegend, eBioscience, or BD unless otherwise indicated (clone in parentheses): B220 (RA3-6B2), CD4 (GK1.5), CD8α (53-6.7), CD11b, CD11b, CD19 (6D5), CD21 (7G6), CD23 (B3B4), CD43 (S7), IgD (11-26c.2a), IgM (R6-60.2), IgMa (DS-1), IgMb (AF6-78) Igκ (187.1), F4/80 (BM8), NK1.1 (PK136), Ter119 (TER-119), goat anti-human Igκ (Southern Biotech), goat anti-mouse IgA (Southern Biotech), goat anti-mouse Igλ (Southern Biotech). Samples were run on a BD LSRII Cytometer.

### Cell Sorting and Adoptive Transfer

Indicated populations of peritoneal B cells from multiple CD45.1-congenic wild type mice were sorted on a FACSAria II cell sorter (Becton Dickinson). After post-sort purity analysis, 1e5 cells were transferred IP into adult (8-12 week) C57BL/6J mice. Recipient mice were analyzed one week following transfer.

### Non-Ig construct cloning and genotyping

Targeting constructs for knock-ins other than Speed-Ig were prepared in largely the same manner. Homology arms exactly flanking the guide site were amplified from genomic DNA of the mouse strain used as embryo donors for microinjection (typically C57BL/6J). A second cloning PCR was performed to provide Gibson overlaps to homology arms and cargo if necessary. Homology arms and cargo DNA were assembled into generic plasmids (pUC18 or pUC19) via Gibson assembly. Microinjections were carried out identically to Speed-Ig, but with crRNAs specific for the relevant guide site. Genotyping knock-in mice was performed by three separate PCRs on DNA samples from potential founder animals. First, PCRs to detect the cargo were performed to identify all mice which carried non-native DNA (e.g. EGFP). Then, cargo^+^ animals were genotyped for targeting PCR across the 5’ and 3’ homology arms to detect properly targeted insertions/replacements. Amplicons arising from targeting reactions were Sanger sequenced to confirm correctly targeted alleles.

## RESULTS

### Ig gene targeting utilizing a Cas9-based approach

To maximize the biological relevance of our Ig knock-in approach, we had the following objectives: (1) to remove the endogenous J segment loci, thereby preventing secondary Ig rearrangements after successful integrations; (2) to use fixed promoter regions which would allow for phenotypic comparisons across mouse lines in a truly Ig-dependent manner; and (3) to minimize targeting construct size and synthesis time such that panels of Ig knock-in mice could be generated quickly and efficiently.

To insert rearranged Ig genes into their corresponding loci, we first selected one guide RNA site each for J_H_1 and J_κ_1, directly upstream of the gene segment coding regions. We then generated targeting constructs with short (∼800bp) “proximal” and long (∼2.2kb) “distal” homology arms bookending all endogenous J segments (**Fig.1A**). Using a single Cas9 guide site, only the proximal homology arm directly abutted the double-strand break and served as an anchor for the intended synthesis-dependent strand annealing. The intervening genomic DNA between the guide RNA site and the beginning of the distal homology arm had no complementarity present in the donor DNA and was excised upon construct integration. Therefore, upon successful integration of the targeting construct, the entire J segment clusters were deleted, preventing secondary Ig rearrangements on the targeted allele. The distal homology arms were selectively increased in length as longer distal, but not proximal, homology arms were demonstrated to increase efficiency of large gene replacements *in vitro*.^43^ Promoter regions 800-1200 base pairs in size including split leader peptide sequences were cloned from the V_H_1 family for heavy chain constructs and the V_κ_10 family for kappa constructs based on usage patterns in wild type B cell repertoires.^34^ Rearranged Ig chains amplified from cDNA of single sorted B cells were assembled seamlessly with promoters into respective targeting constructs using Gibson assembly^40^ (**Supplementary Fig.1**). This modular construct assembly allowed for rapid and scalable Ig cloning combined with flexibility in promoter choice. Fertilized murine oocytes were microinjected at the one-cell stage with targeting constructs and either single-guide RNA (sgRNA) with Cas9 mRNA or pre-complexed gRNA:Cas9 ribonucleoproteins (RNPs) (**Fig.1B**). Potential Ig knock-in founders were screened for intact Ig insertions via ‘targeting’ polymerase chain reaction (PCR) which was accommodated by homology arms much shorter than those used in traditional knock-in transgenesis (**Fig.1C**). Ig inserts were further verified by southern blotting (**Fig.1D**). Rarely, some founder animals possessed off-target integrations (**Supplementary Fig.2**). After several rounds of optimization, successful Ig integrations/J cluster deletions ultimately reached 20.7% of live-born mice for heavy chains and 23.3% for kappa chains (**Table 1**). Successful knock-ins were only recovered from the injection of targeting constructs with longer 3’ homology arms (**Table 1**). Notably, the use of gRNA:Cas9 RNPs was superior to sgRNA/Cas9 mRNA, similar to recent reports.^41,44^

**Figure 1.**
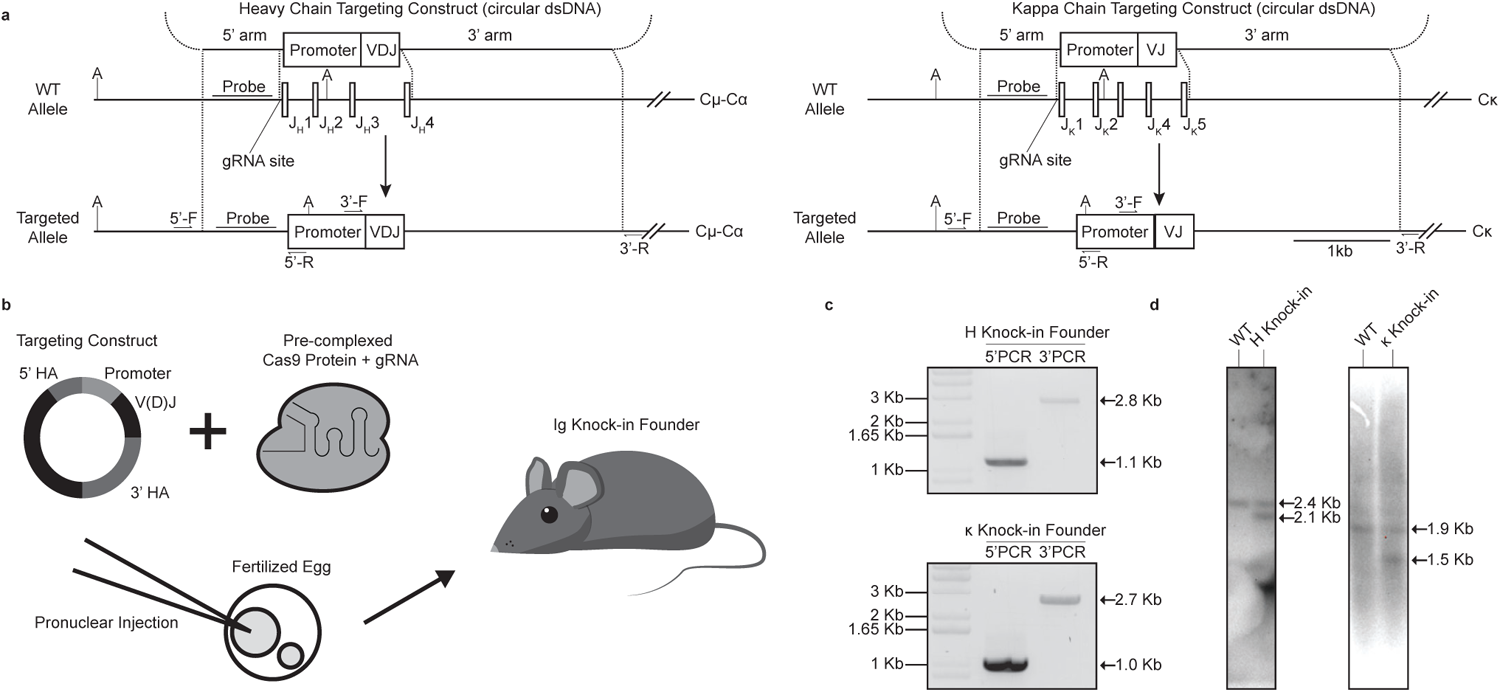
Targeted integration of rearranged V regions at the J_H/_J_κ_ loci. **A)** Targeting approach to replace endogenous J segments with rearranged VDJ_H_/VJ_κ_. Dashed lines indicate areas of complementarity for homology arms. **B)** Schematic diagram of pronuclear injection **C)** Genotyping of H and κ knock-in founders using paired targeting PCRs. Corresponding primers are depicted in panel A. **D)** Southern blotting of H and κ knock-in mice. Genomic DNA was digested with ApoI (for H) or AccI (for κ) and probed within the 5’ homology arm. A: ApoI/AccI sites.

**Table 1.**
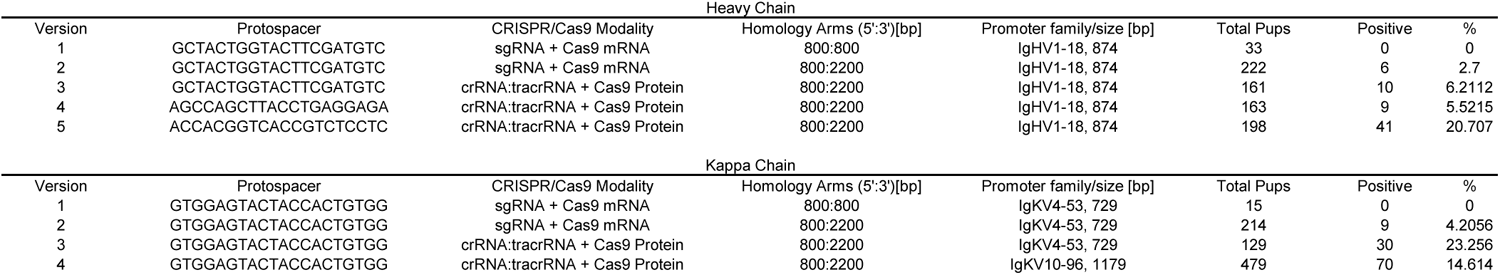
Speed-Ig knock-in efficiencies.

### Speed Ig expression, allelic exclusion, and BCR-driven phenotypes

We performed a series of experiments to evaluate the utility of Speed-Ig mice in studying B cell biology, a hallmark of prior Ig knock-in approaches. Heavy chain and light chain expression were measured independently by flow cytometry. Progeny from H knock-in mice (on the *IgH*^*b*^ C57BL/6J background) bred to *IgH*^*a*^ allotype mice demonstrated knock-in chain expression at the protein level (**Fig.2A**, left panels). As expected, the knocked-in chain, due to its rearranged state and ensuant expression kinetics, prevented endogenous rearrangements and therefore excluded *IgH*^*a*^ allele expression^45^. On an *IgH*^*Δ*^ background, knocked-in heavy chains were capable of driving normal B cell development (**Supplementary Fig.3A**). Furthermore, the surface expression of IgM and IgD were comparable between wild type B cells and B cells from *IgH*^*KI/Δ*^ mice (**Supplementary Fig.3A**). The *IgH*^*Δ*^ background also provided an opportunity to assess class-switch recombination in Speed Ig mice. As expected, B cells forced to use knocked-in H chains were able to switch to the IgA isotype in mucosal tissue, a firm indication that our Speed-Ig lines retain this important capability (**Supplementary Fig.3B**). These data confirm the functionality of the targeting approach, promoter regions, and Ig integrations for H knock-ins. Similar experiments were performed using κ knock-in mice. κ knock-ins were bred to mice possessing the human kappa constant region to allow for allele-specific flow cytometry staining.^31^ κ knock-in mice with V_κ_10-96 promoters had normal mIgκ surface expression and outcompeted the human kappa allele (**Fig.2A**, right panels). This contrasted with κ knock-in mice generated with V_κ_4-53 promoter regions, which had poor kappa expression (**Supplementary Fig.4**). Although the V_κ_4-53 promoter region was selected based on published work^46^, we did not observe adequate Igκ expression in our lines, prompting our use of a larger promoter region from a different Igκ family. We then examined full (H + κ) Ig knock-in mice (henceforth with V_κ_10-96 promoters) for B cell developmental progression. It has been previously noted that, in Ig knock-in mice, developing B cells transit the pre-B stage faster than wild-type cells due to pre-rearranged Ig expression.^9,46–48^ Our knock-in lines demonstrated a reduction in pre-B frequency, consistent with prior knock-in approaches and further validation of knocked-in Ig expression (**Fig.2B**). Taken together, these results suggest Speed-Ig mice display physiologic Ig chain expression levels and that knocked-in Igs predominate over endogenous chains. In summary, our results demonstrated that the Speed-Ig system is a robust tool for *in vivo* studies of monoclonal B cell biology.

**Figure 2.**
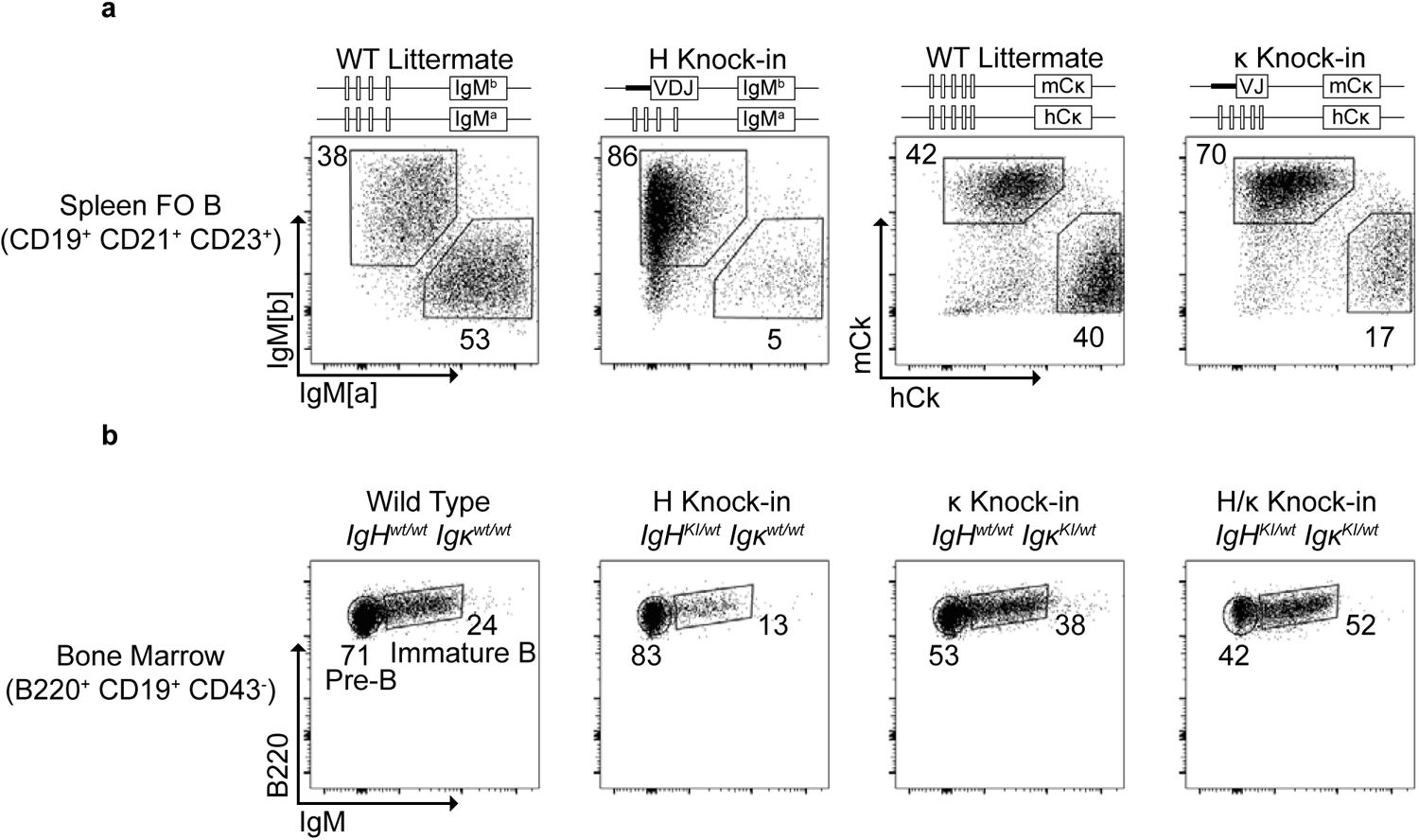
Speed-Ig gene expression and B cell development. **A)** Knocked-in H and κ chains dominate over endogenous alleles. [Left panels] Flow cytometry of splenic follicular B cells (CD19^+^ CD21^+^ CD23^+^) from *IgH*^b/a^ and *IgH*^KI/a^ littermates stained with allotype-specific anti-IgM antibodies. *IgH* locus genotype indicated above plots. [Right panels] Flow cytometry of splenic follicular B cells from *Igκ*^m/h^ and *Igκ*^KI/h^ littermates stained with human and mouse-specific anti-IgK antibodies. *Igκ* locus genotype indicated above plots. **B)** Bone marrow development of Speed-Ig B cells expressing H, k, or both Ig knock-in chains. All panels are representative of n ≥ 3 mice.

### Peritoneal B cell relationships in Ig knock-in mice

Having validated our knock-in system, we sought to understand the ontogeny of the less well-studied innate-like B cell populations: non-canonical B-1a and B-1b cells. To do so, we created mouse lines with Ig pairs from randomly selected B-1b clones or non-expanded B-1a clones (**Table 2**)^34^. Additionally, we created a canonical B-1a V_H_11-V_κ_9 Ig line “321C6” to serve as a control for BCR-driven cell fating^25,49^. Both canonical and non-canonical B-1a Ig knock-in mice harbored populations of B cells strongly biased towards the B-1a fate (**Fig.3A-B**). B-1b Ig knock-in mice, on the other hand, gave rise to both B-1 and B-2 cell subsets in the peritoneal cavity, albeit with a reduction in B-1a (**Fig.3A-B**). The same result was observed in B-2 Ig knock-in lines, such that B-1b and B-2 Ig knock-in lines were phenotypically indistinguishable. Given the possibility that some endogenous early-life B-1a precursors escaped allelic exclusion, we bred one B-1a line, “416C1, and B-1b line, “301G2”, to *Rag1*-deficient backgrounds. In this setting, virtually all peritoneal B cells in 416C1/*Rag1*^*-/-*^ mice were of the B-1a phenotype as expected (**Fig.3C**). Conversely, 301G2/*Rag1*^*-/-*^ mice had minimal B-1a cells, confirming that the knocked-in Ig pair was incapable of giving rise to the B-1a subset (**Fig.3C**). However, there was no loss of peritoneal cavity B-2 cells, suggesting that some Ig pairs can seed both B-1b and B-2 subsets. Interestingly, ∼15% of the B cells present in B-1b and B-2 Ig knock-in lines did not fully conform to either the B-1 or B-2 lineages, appearing as an “intermediate” (Int) cells. As Int cells persisted in the *Rag1*-deficient 301G2 line, we hypothesized that shared origins between the B-1b and B-2 subsets might exist in wild type animals. However, due to the lack of markers used to identify B-1b cells, it was possible that immature or transitional B cells were present in our samples. To rule out the presence of young CD23^variable^ B cells which might appear in the Int gate, we utilized the *Rag2-EGFP* reporter line. In these animals, ∼80% of splenic transitional stage 1 (T1) and stage 2 (T2) cells still had residual GFP signal, indicative of recent development (**Supplementary Fig.5A**). Splenic follicular B cells (FO B) and marginal zone B (MZ B) cells were largely GFP^-^, in concordance with their temporal separation from lymphopoiesis (**Supplementary Fig.5A**). Virtually all cells in the peritoneal cavity were GFP^-^, suggesting that the large fraction of poorly defined Int B cells present are not recent bone marrow emigrants (**Supplementary Fig.5A**). As another determination of lineage relationships, we pursued a fate-mapping experiment by crossing *CD23-Cre* with *R26R-EYFP* mice. CD23 expression initiates at the T2 stage of development and is ubiquitously expressed by B-2 cells. In the peritoneal cavity, approximately 98% of B-2 phenotype cells were fate-mapped, demonstrating efficient and robust Cre activity driven by the CD23 promoter (**Supplementary Fig. 5B**). Nearly 90% of Int cells were also YFP^+^, in line with the variable but apparent surface CD23 expression. Notably, ∼70% of B-1b cells were also fate-mapped, suggesting prominent seeding of the B-1b compartment by cells that had expressed CD23 during a precursor stage (**Supplementary Fig.5B**). B-1a cells displayed the lowest fate-mapping by CD23, with approximately 30% YFP^+^ (**Supplementary Fig.5B**). Taken together, these data suggest that some fraction of each B-1 lineage has origins in mature, CD23^+^ precursors. Furthermore, the ability of a single Ig pair to drive B-1b, B-2, and intermediate cell phenotypes establishes the possibility of shared B cell lineages in the peritoneal cavity.

**Table 2.**
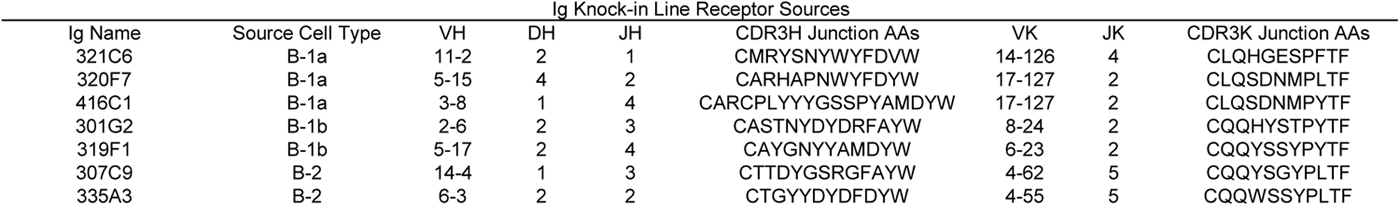
Ig pairs used for knock-in lines.

**Figure 3.**
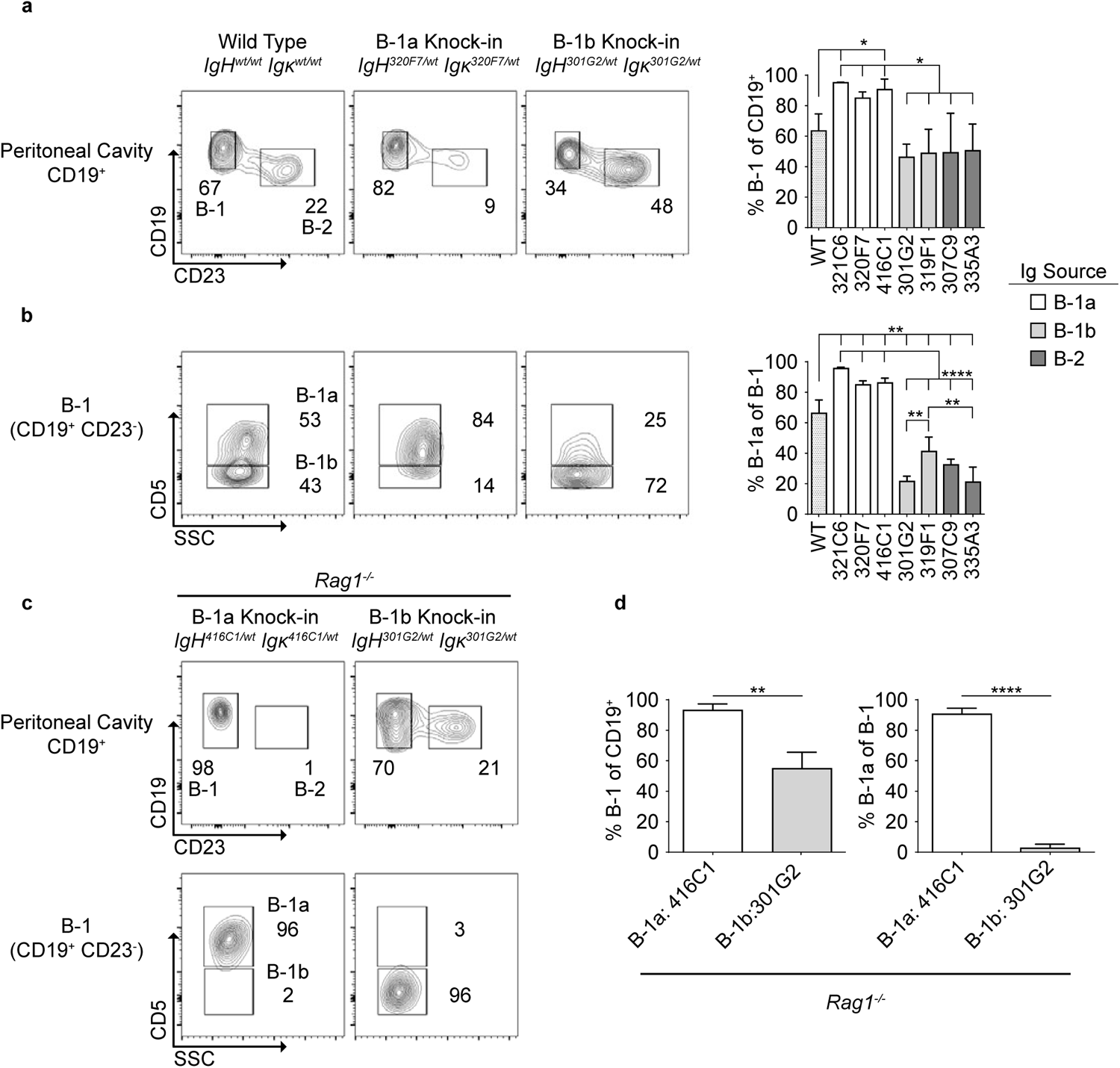
B-1b and B-2 Ig knock-ins display unpredicted phenotypic flexibility. **A)** [Left] Representative plots of peritoneal B cells in wild type, non-canonical B-1a, and B-1b Ig knock-in mice. [Right] Frequencies of B-1 cells of all peritoneal B cells in Ig knock-in lines. Significance determined by one-way ANOVA with post-hoc Tukey test. **B)** [Left] Breakdown of B-1a and B-1b cells present in the B-1 gate of representative Ig knock-in lines as in A. [Right] Frequencies of B-1a and B-1b cells in all Ig knock-in lines. Significance determined by one-way ANOVA with post-hoc Tukey test. **C)** Comparison of a non-canonical B-1a line and a B-1b line on Rag1-defecient backgrounds. **D)** frequency of B-1 among all B cells and B-1a among B-1 cells in Rag1-deficient Ig knock-in lines. Significance determined by Student’s T-test. n≥ 3 biological replicates for all panels.

### *Zbtb32* and *Bhlhe41* identify B-1-like precursors in the peritoneal cavity

To further analyze the ontogeny of B-1 cells, we created a *Zbtb32* transcription factor reporter line (**Supplementary Fig.6A**). Zbtb32 is highly upregulated in B-1 cells, but has minimal expression in follicular B cells^17,18^. As expected, virtually all peritoneal B-1 cells were EGFP^+^ (**Fig.4A-B**). Interestingly, Int cells were largely positive for EGFP expression and a small fraction of B-2 also had expression (**Fig.4A-B**). The mean fluorescence intensity of GFP across the three populations suggests a gradual phenotypic transition from B-2 to B-1 (**Fig.4B**, right panel). We then compared B-2 cells grouped by the presence or absence of EGFP expression. Across a variety of markers and cell size and granularity, EGFP^+^ B-2 cells were skewed toward a B-1 phenotype compared to GFP^-^ B-2 cells (**Fig.4C**). Zbtb32 expression therefore directly correlates with B-1 phenotype polarization, and putative B-1 precursors present within the B-2 subset can be identified by Zbtb32 expression.

**Figure 4.**
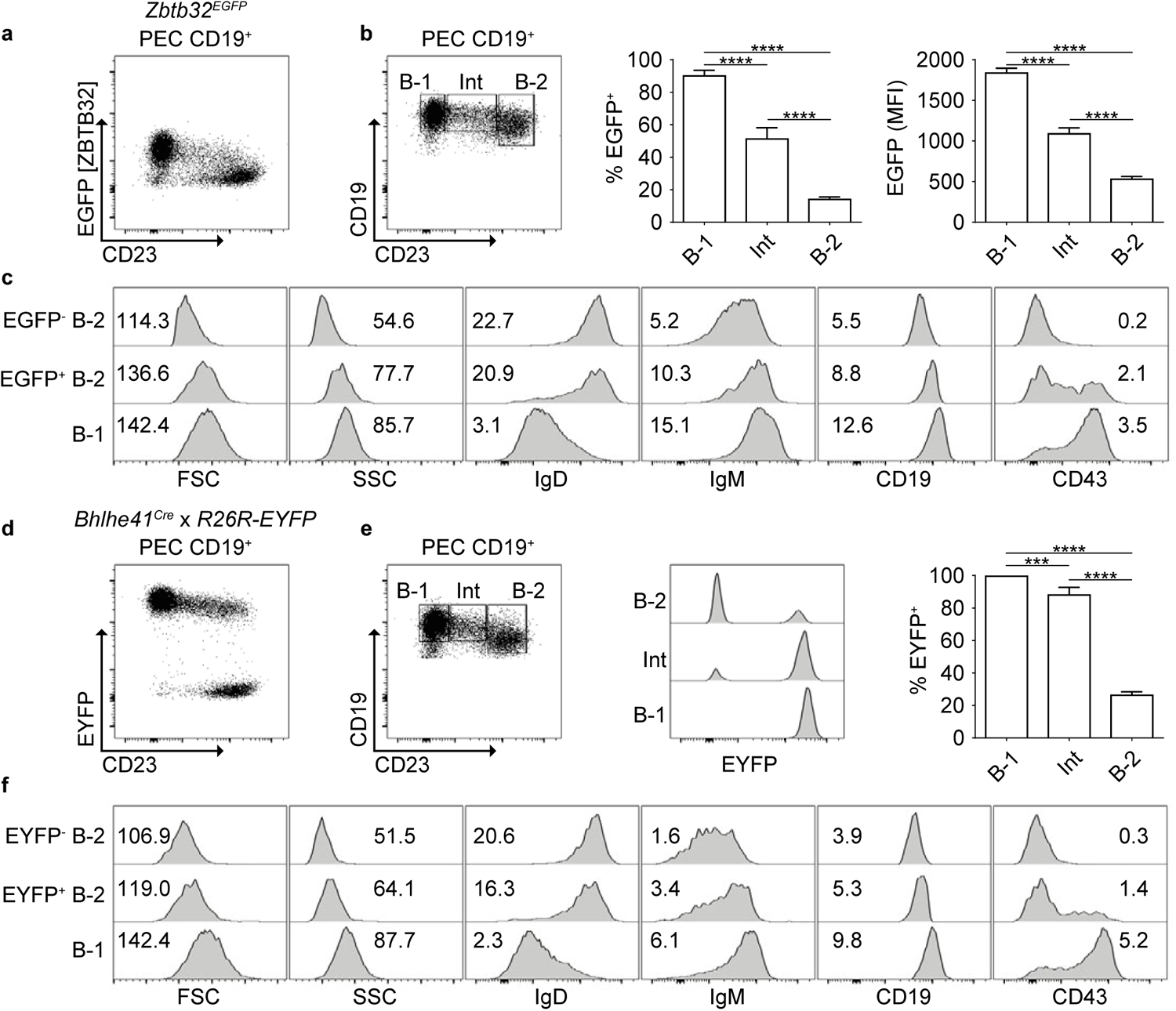
A *Zbtb32* reporter and *Bhlhe41* Cre-driver reveal B-1-like phenotypic changes in peritoneal B cells. **A)** Zbtb32 expression reported by EGFP in peritoneal B cells. **B)** EGFP^+^ frequency and mean fluorescence intensity in peritoneal B cell subsets. Significance determined by one-way ANOVA with post-hoc Tukey test. **C)** Histograms of scatter values and fluorescence intensities for indicated cell subsets. Mean fluorescence intensities given as 1×10^3^. **D)** Fate-mapping of peritoneal B cells in *Bhlhe41*^*Cre*^ x *R26R-EYFP* mice. **E)** EYFP^+^ frequency in peritoneal B cell subsets. Significance determined by one-way ANOVA with post-hoc Tukey test. **F)** Histograms of scatter values and fluorescence intensities for indicated cell subsets. Mean fluorescence intensities given as 1×10^3^. n≥ 4 biological replicates for all panels.

We further probed peritoneal B cell lineage relationships by generating a *Bhlhe41*^*Cre*^ line (**Supplementary Fig.6B**). *Bhlhe41* is a master factor in B-1 differentiation and like *Zbtb32* is highly expressed in B-1 cells with negligible expression in other B cell subsets^16–18^. Like *CD23*^*Cre*^, we crossed the *Bhlhe41*^*Cre*^ line to *R26R-EYFP* mice to delineate the ontogeny of B-1 and Int cells. We observed that all B-1 were fate-mapped by *Bhlhe41*^*Cre*^, as were ∼90% of Int cells (**Fig.4D-E**) Consistent with the Zbtb32 reporter data, a fraction of peritoneal B-2 cells were fate-mapped by Bhlhe41Cre, providing a second line of evidence for transcriptional heterogeneity of the peritoneal B-2 compartment (**Fig.4D-E**). Gating fate-mapped peritoneal B-2 cells from EYFP^-^ B-2 cells confirmed that fate-mapped B-2 appear more B-1-like across a variety of markers in addition to cell size and granularity (**Fig.4F**). These data were consistent with observations in the *Zbtb32* reporter line. Taken together with results from Speed-Ig lines, we hypothesized some peritoneal B-2 cells were triggered to differentiate to a B-1 phenotype, and that Int cells represent ex-B-2 cells en route to the B-1 cell fate.

### Peritoneal Cavity B-2 and Int cells seed the B-1 compartment

To test the ability of peritoneal B-2 and Int cells to see the B-1 compartment, we performed an adoptive transfer experiment using congenically marked but otherwise wild type B cells (**Fig.5A**). Cells were FACS-purified and assessed for purity prior to intraperitoneal injection into C57BL/6J hosts (**Fig.5A-B**). After one week, cells were collected from recipient animals by peritoneal lavage and for stained for B cell makers. Both B-1a and B-1b subsets largely retained their original phenotypes following transfer, consistent with a previous report^50^ (**Fig.5B-C**).Transferred Int cells closely resembled the phenotype of transferred B-1 cells, with most cells now falling into a CD23^lo/-^ gate (**Fig.5B-C**). A large fraction of transferred B-2 cells also fell into the B-1 gate at a frequency in line with the *Zbtb32*^*EGFP*^ and *Bhlhe41*^*Cre*^ results (**Fig.5.B-C**). The presence of host B cells allowed us to directly compare the marker profiles of donor cells. Not only had Int and B-2 cells now appearing in the B-1 gate lost surface expression of CD23, they had gained a marker profile consistent with the B-1 phenotype (**Fig.5D**). Regardless of original cell phenotype, the donor cells falling into a B-1 gate were indistinguishable from bona fide host B-1 cells, allowing us to conclude that some B-2 and most Int cells can differentiate to the B-1 fate. In summary, the Speed-Ig knock-in lines, transcription factor genetic approaches, and adoptive transfers all suggest an unappreciated shared origin of peritoneal B cell subsets, with previous instructive and induced-differentiation models only accounting for B-1a cells.

**Figure 5.**
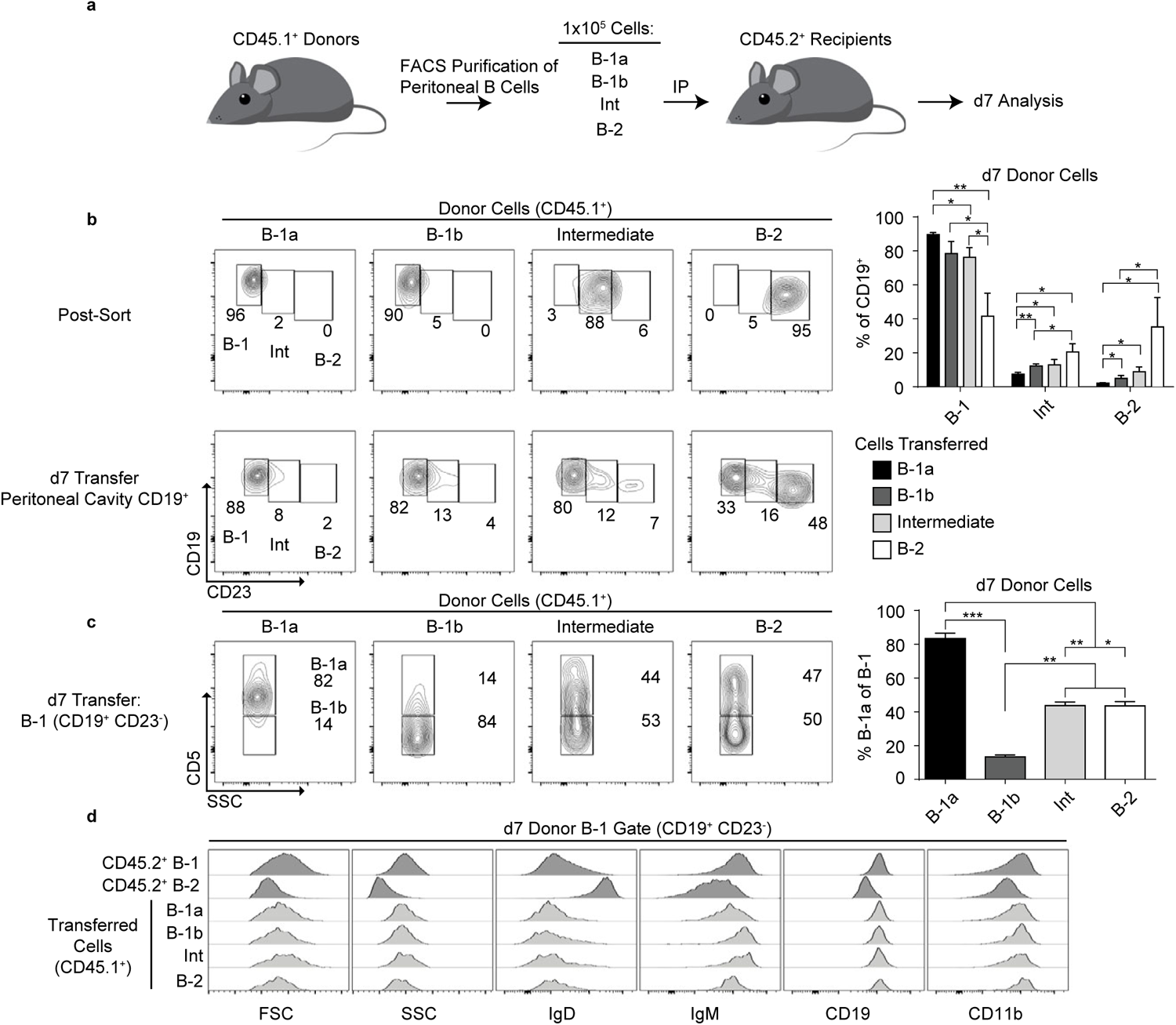
Adoptively transferred peritoneal B-2 and Intermediate cells give rise to B-1 cells. **A)** adoptive transfer scheme. **B)** Post-sort and d7 analysis of donor cells. Significance determined by Student’s T-test. **C)** Breakdown of B-1a and B-1b subsets within the B-1 gate of donor cells. Significance determined by one-way ANOVA with post-hoc Tukey test. **D)** Histograms of scatter and mean fluorescence intensities for donor cells present in the B-1 gate compared to host B-1 and B-2 cells. n≥ 3 biological replicates for all panels.

## DISCUSSION

Here, we reported an efficient and robust Cas9-mediated method to generate Ig knock-in mice. Our method retains all the biologically useful features of targeted Ig mouse lines while increasing throughput and reducing generation time. Knocked-in Igs expressed at levels similar to wild-type mice, strongly excluded endogenous rearrangements, and recapitulated physiologically relevant BCR signaling. Using identical promoter regions across Speed-Ig lines provides several advantages over using V region-specific promoters. The promoters themselves are now well validated to provide robust expression. Phenotypic comparisons between Speed-Ig lines can be directly attributed to different V regions, as only these will differ among lines.

Our approach complements the elegant method recently reported by Victora and colleagues^11^. Their strategy effectively halves the number of mouse lines required to study any given Ig pair by virtue of integrating both Ig chains into the heavy chain locus. Such mouse lines are a valuable tool to study mature B cell responses to cognate antigens. Our method, while still requiring separate integration events at the *IgH* and *Igκ* loci, represents a closer approximation of endogenous Ig expression kinetics and amplitude. Depending on the biological question, each strategy has clear advantages and disadvantages. Our heavy chain knock-in approach is similar to that of Batista and colleagues; Lin et al. demonstrated impressive integration efficiencies at the *IgH* locus^10^. The main difference between our approaches is the length of targeting construct homology arms. Increasing the homology arms to lengths commonly used for embryonic stem cell constructs might afford the greatest efficiency. Our chief comparative advantage is the ability to genotype via targeting PCR across both homology arms to ensure correct integrations, in addition to our description of *Igκ* knock-ins. Nevertheless, there is clearly a relationship between cargo size and homology arm lengths that affects integration efficiency. The Speed-Ig strategy stands to be improved by further studies into optimal homology arm length, especially for the longer distal arm. Guide site selection was originally conducted by analyzing loci with CHOPCHOP and the Broad Institute’s sgRNA Designer^51–53^. These algorithms are based off screening results of sgRNA libraries in cell lines. Later guide site selection was through the IDT design tool for crRNA:tracrRNA bipartite gRNA which may behave differently from sgRNA *in vivo*^44^. While all targeted integrations were achieved regardless of predicted score, our highest efficiencies came from guides with on-target scores above 60 per IDT. However, guide site quality is only one factor in determining knock-in rates, as the length of inserted cargo can also have an effect^54^. Our results are informative on the insertion of cargo up to at least two kilobases. This range includes most if not all reporter genes and a variety of genetic tools, including Cre recombinase and luciferases. In conclusion, our method may serve as a platform for universally successful gene editing in the mouse.

Our novel insights into B-1 plasticity principally arose from observations in Speed-Ig mice. The ability of a single Ig pair to drive cells into the B-2 and B-1b subsets Recent evidence for lineage plasticity in vivo using an inducible Ig expression system demonstrated the ability of B-2 cells to acquire a B-1 phenotype^18^. While elegant, this approach was still limited to the biology of canonical B-1a signaling and differentiation. In a related study, BCR signaling blockade prevented B-1a Ig transgenic cells from assuming a full B-1a phenotype^28^. The apparent heterogeneity of peritoneal B cell subsets limits extrapolation of these findings. We expanded on existing data by showing that a monoclonal knocked-in Ig pair can give rise to diverse B cell subsets, and that a phenotypic spectrum is apparent in wild type mice. The polarity of cells toward the B-1 phenotype was directly correlated with the expression of hallmark B-1 transcription factors, regardless of CD23 surface expression. Furthermore, we observed rapid differentiation of peritoneal B-2 and Int cells into B-1 cells following adoptive transfer. Similar results were observed in an adoptive transfer experiment using peritoneal B-2 cells, but not splenic B-2 cells^55^. These data suggest that BCR-independent signaling events occurring in the peritoneum might drive acquisition of the B-1 phenotype in a broad range of B cells. This argument is supported by the fact that peritoneal B-2 cells have no repertoire bias^21^. The continual presence of Int cells in wild type mice opens the possibility that at steady-state, there is perpetual seeding of the B-1 compartment by non-B-1 precursors. Moreover, in combination with data from B-1b and B-2 Ig knock-in mice, there appears to be some factor(s) limiting the size of the B-1 niche. Both BCR-dependent and -independent mechanisms are possible, and further studies are needed to identify the rules governing innate-like B cell fating.

## ACKNOWLEDGEMENTS

We are grateful to H. Zhao, J. Fan and X. Chen of the Transgenic Core at the University of Chicago for technical assistance with embryo microinjections and advice on southern blotting. We thank M. Chowdury and B. Kee for provision of the *Rag2-EGFP* mouse line and D. Kennedy and M. Clark for provision of the *CD23*^*Cre*^ mouse line. We also thank T. Golovkina for helpful discussions.

## Contributions

S.E. conceived the study, designed and performed experiments, analyzed data, and wrote the paper. E.Z. performed experiments, L.D helped generate the knock-in models, A.B. conceived the study and wrote the paper.

---

**Supplementary Figure 1.**
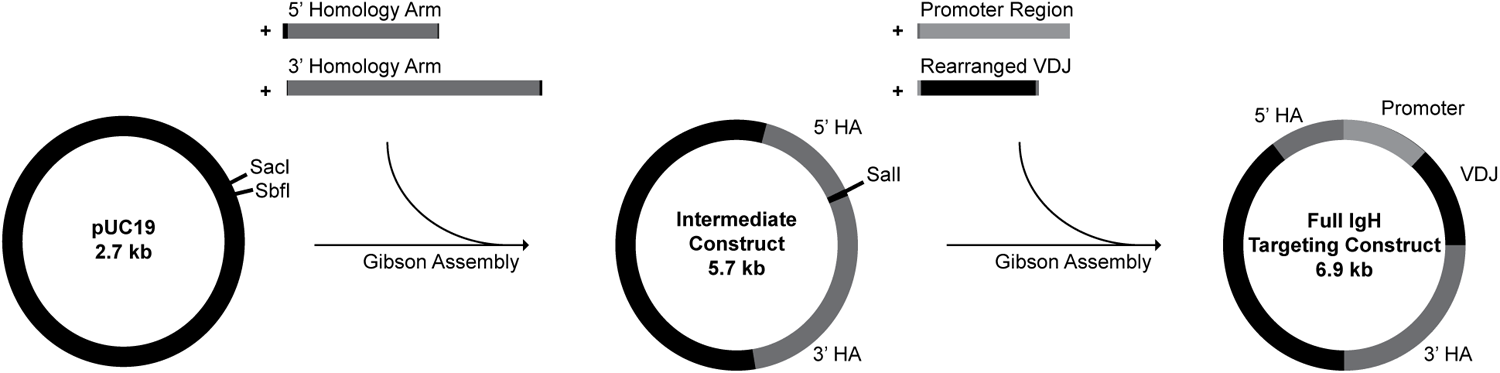
Rapid *IgH* targeting construct generation. A generic plasmid backbone was digested with SacI and SbfI, allowing assembly of homology arms into an intermediate targeting construct. Intermediate constructs were digested with SalI, followed by final assembly with an Ig-specific promoter region and Ig variable regions of interest. Regions of assembly overlaps at the ends of inserts are color-matched to construct sequence. A similar assembly pipeline was done for *Igκ* targeting constructs. Not depicted to scale.

**Supplementary Figure 2.**
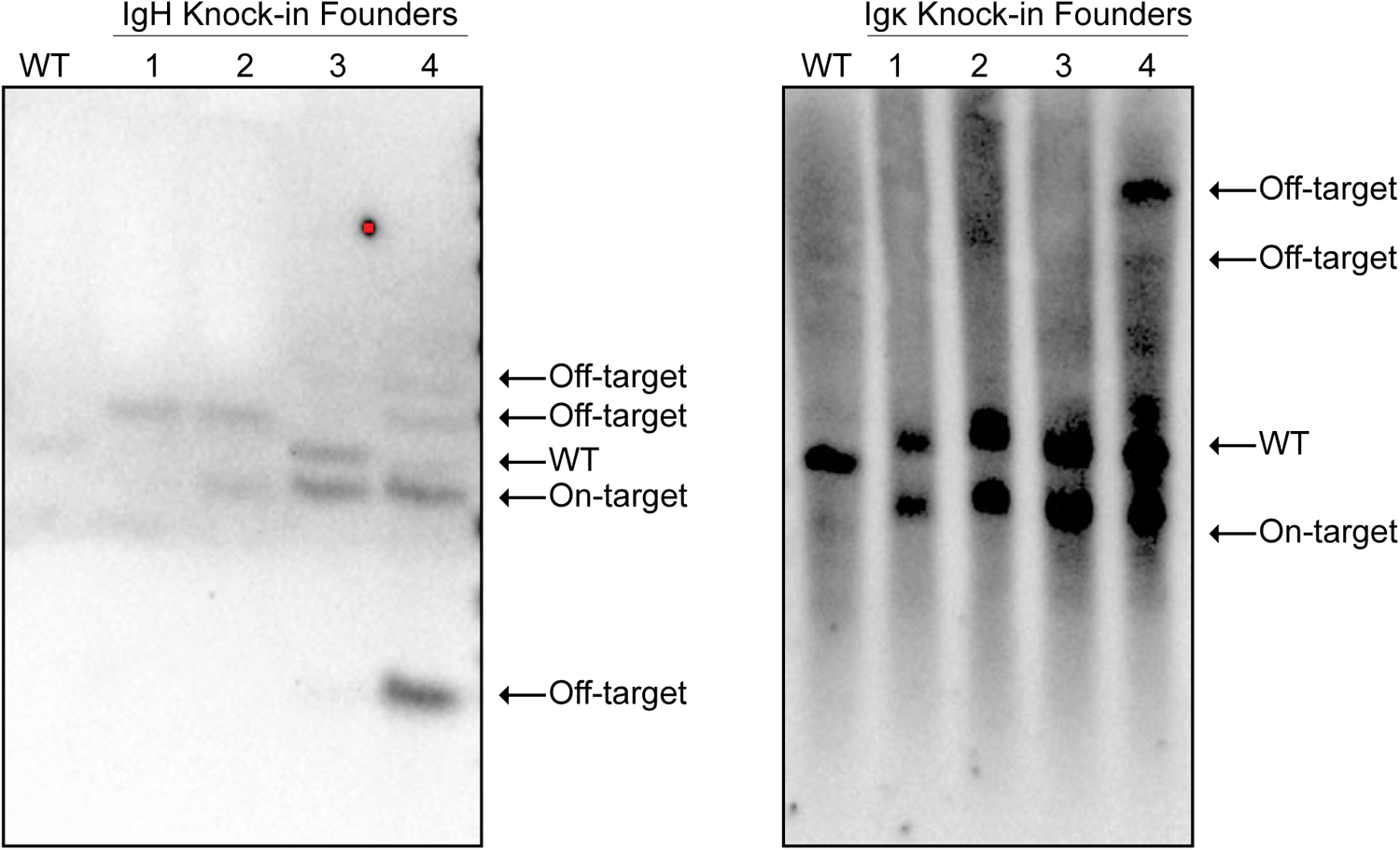
Off-target analysis of H and κ knock-in founder animals. Genomic DNA was digested with ApoI (for H) or AccI (for κ) and probed within the 5’ homology arm. Wild type, on-target, and off-target bands are indicated by arrows. Panels include representatives from four different knock-in lines.

**Supplementary Figure 3.**
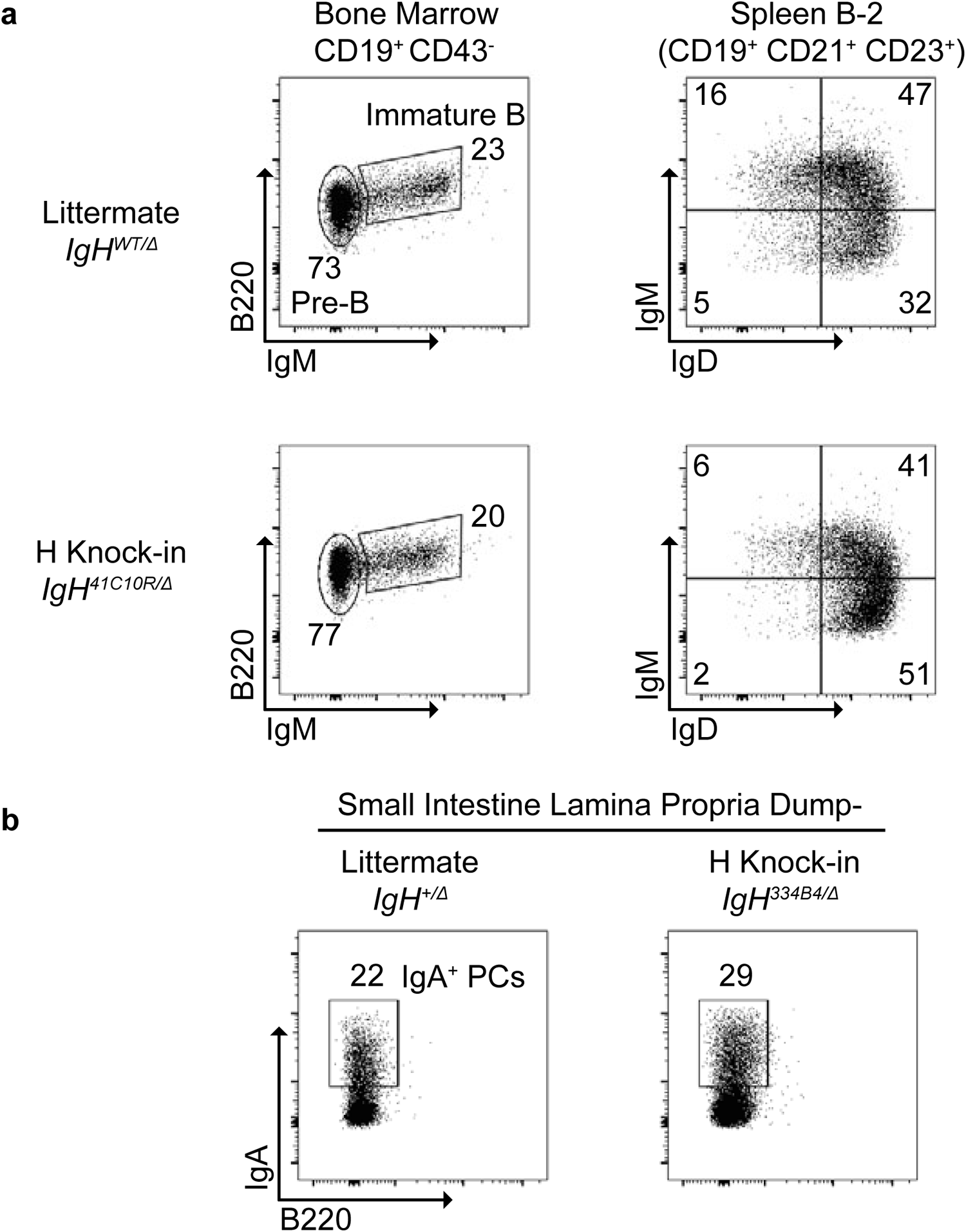
H knock-ins are sufficient in the absence of endogenous alleles and can class switch. **A)** [Left panels] Flow cytometry of developing B cells in the bone marrow and [right panels] mature B cells in the spleen of an H knock-in mouse on an IgH-deficient background. **B)** Class switching to IgA in the gut lamina propria of an *IgH*^*KI/Δ*^ mouse All panels are representative of n ≥ 3 mice.

**Supplementary Figure 4.**
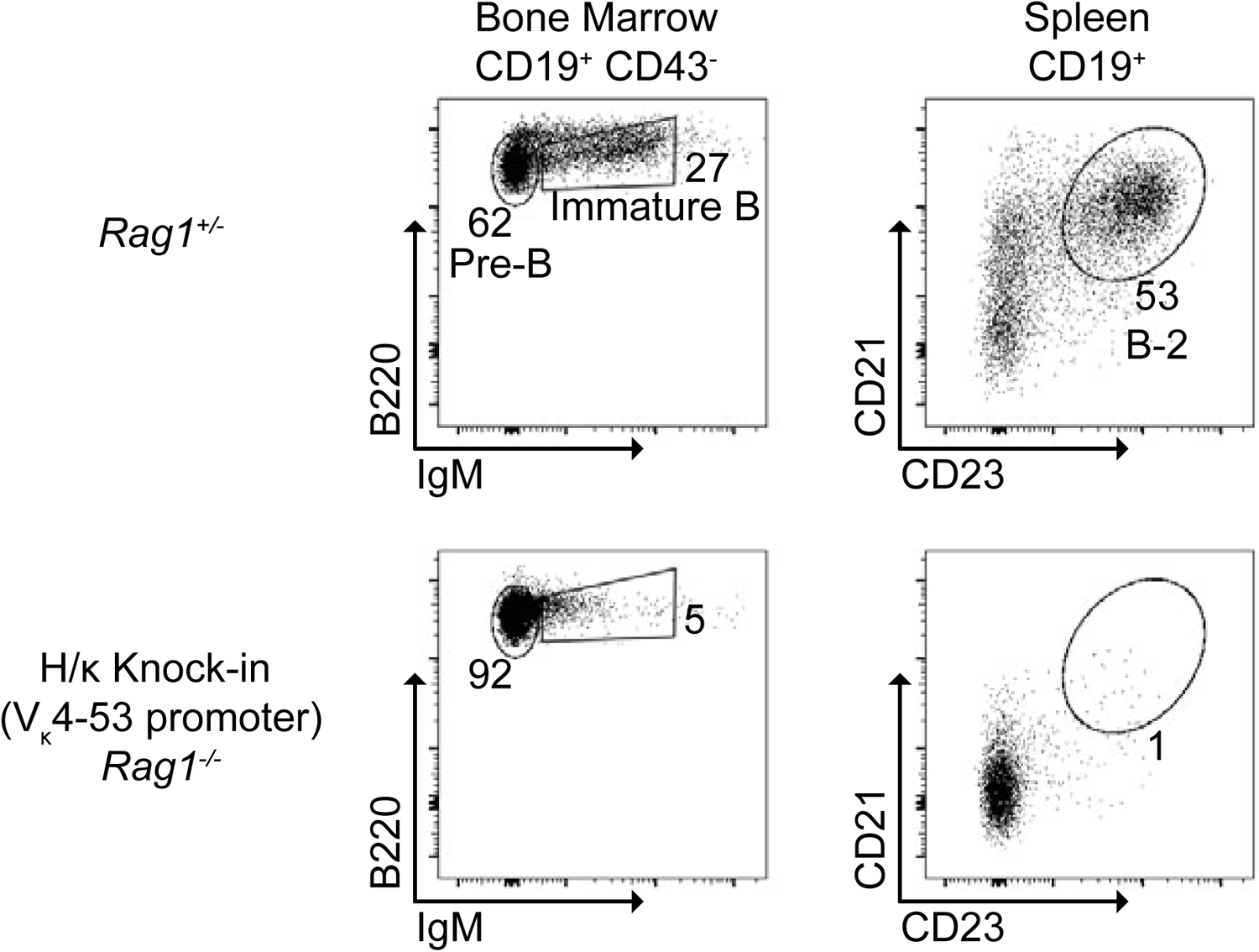
A V_κ_4-53 promoter fragment is insufficient to drive Igκ knock-in expression. [Left panels] B cell development in the bone marrow gated on CD19^+^ cells past the Pro-B (CD43^+^) stage. [Right Panels] surface expression of CD21 and CD23 on splenic CD19^+^ B cells. All panels are representative of n≥ 3 mice from 5 separate lines using the V_κ_4-53 promoter

**Supplementary Figure 5.**
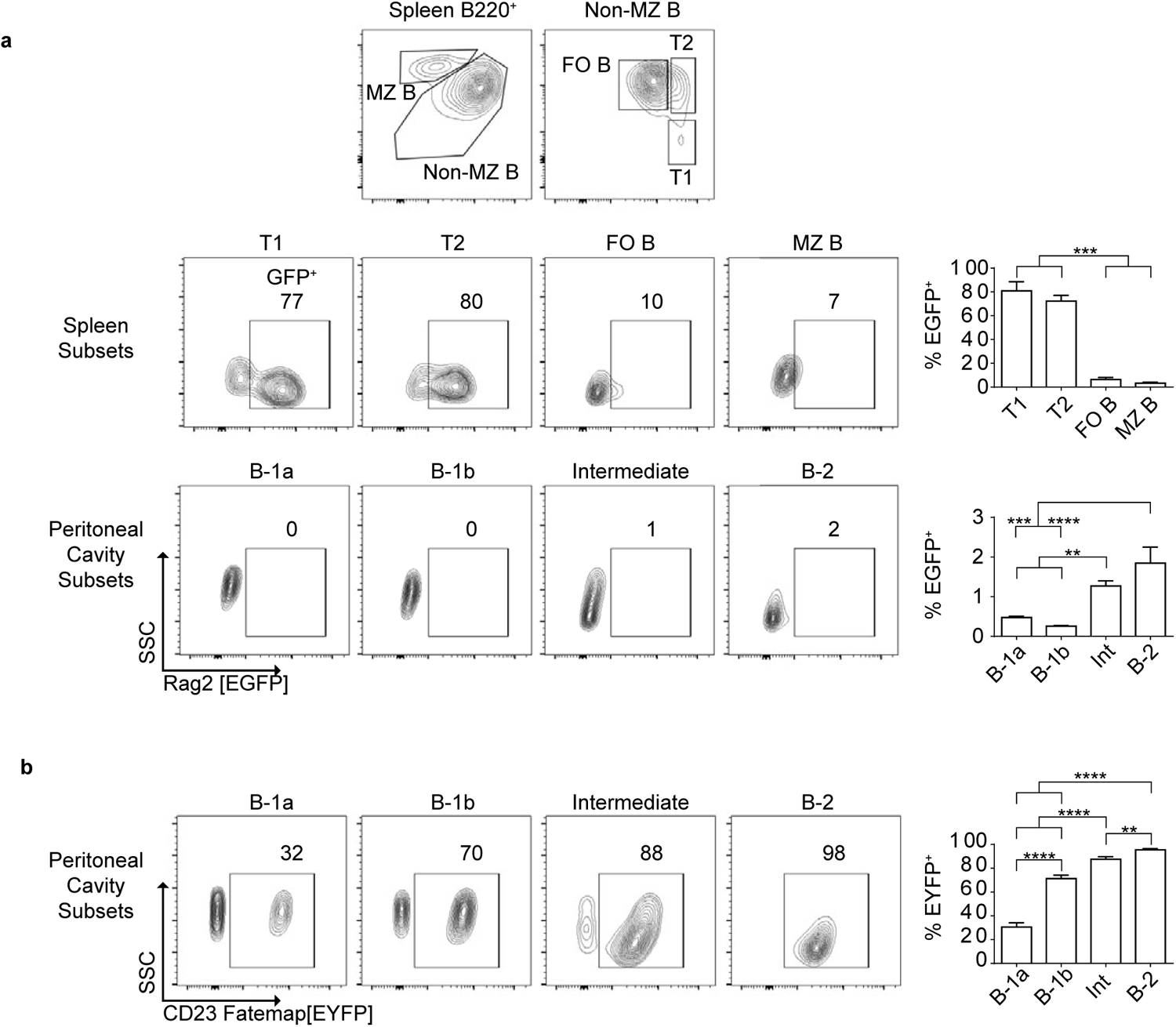
CD23^+^ precursors contribute to B-1 subsets with minimal contribution from recent bone marrow emigrants. **A)** [Top panels] gating strategy for splenic T1, T2, FO B and MZ B subsets in *Rag2-EGFP* mice. [Bottom panels] frequency of GFP^+^ (recent bone marrow emigrants) in splenic and peritoneal B cell subsets. Significance determined by one-way ANOVA with post-hoc Tukey test. **B)** Peritoneal B cells from *CD23-Cre* x *R26R-EYFP* mice. Significance determined by one-way ANOVA with post-hoc Tukey test. n≥ 3 biological replicates for all panels.

**Supplementary Figure 6.**
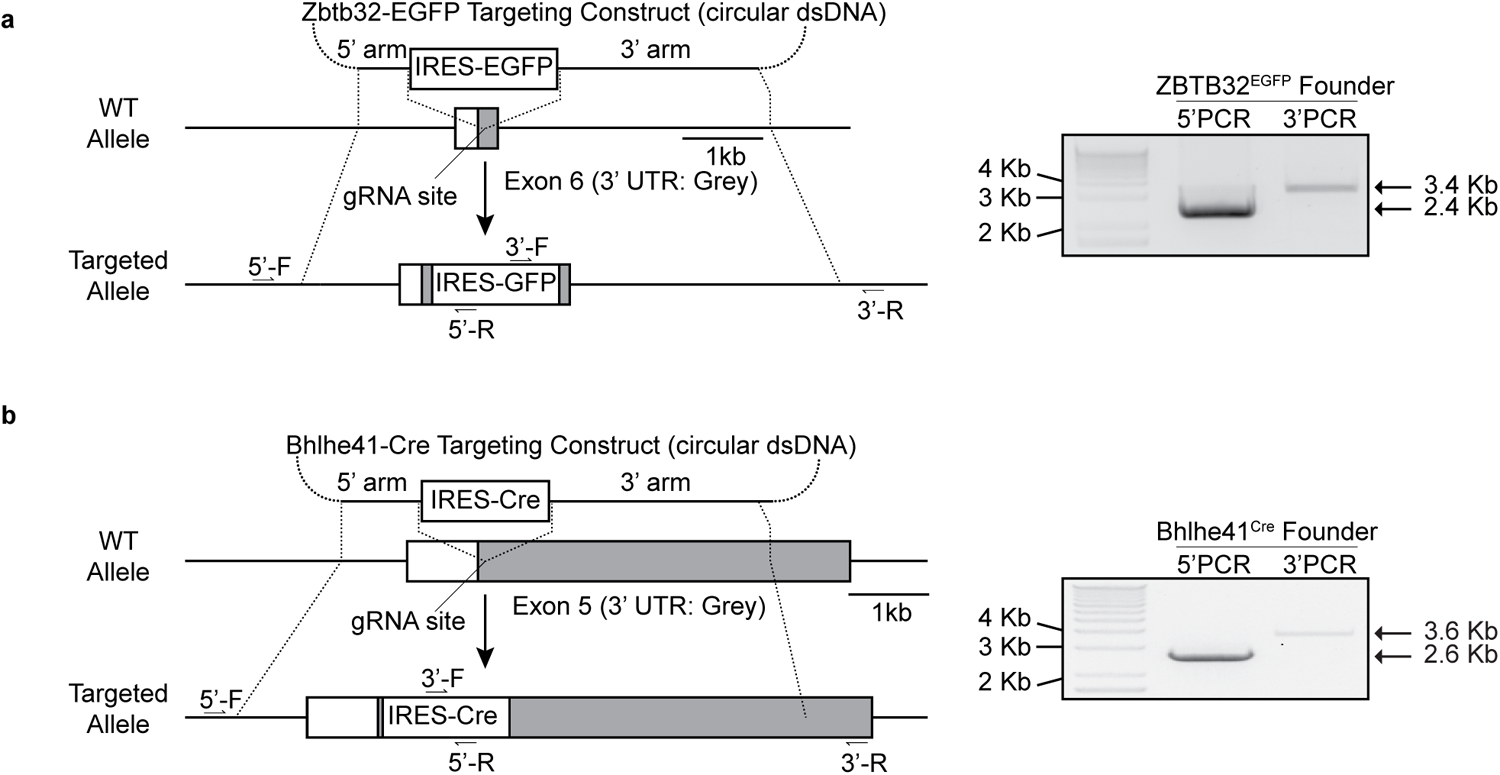
Generation of *Zbtb32*^*EGFP*^ and *Bhlhe41*^*Cre*^ lines. **A)** [Left]Targeting strategy for Cas9-mediated insertion of IRES-led EGFP into the 3’ UTR of the *Zbtb32* locus. [Right] Genotyping PCR bands of a founder animal. **B)** [Left] Targeting strategy for Cas9-mediated insertion of IRES-led Cre into the 3’ UTR of the *Bhlhe41* locus. [Right] Genotyping PCR amplicons of a founder animal.

**Supplementary Table 1.**
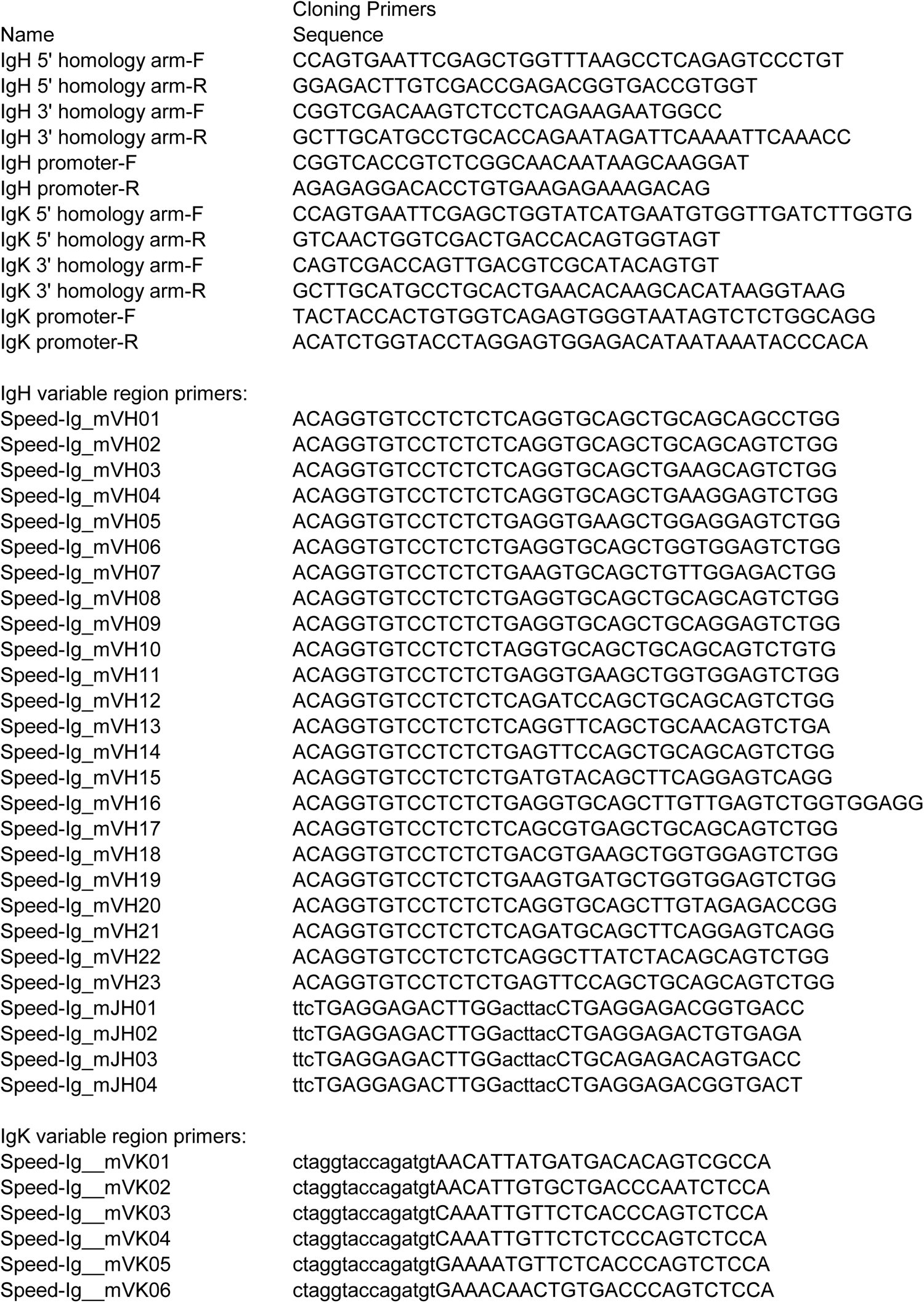
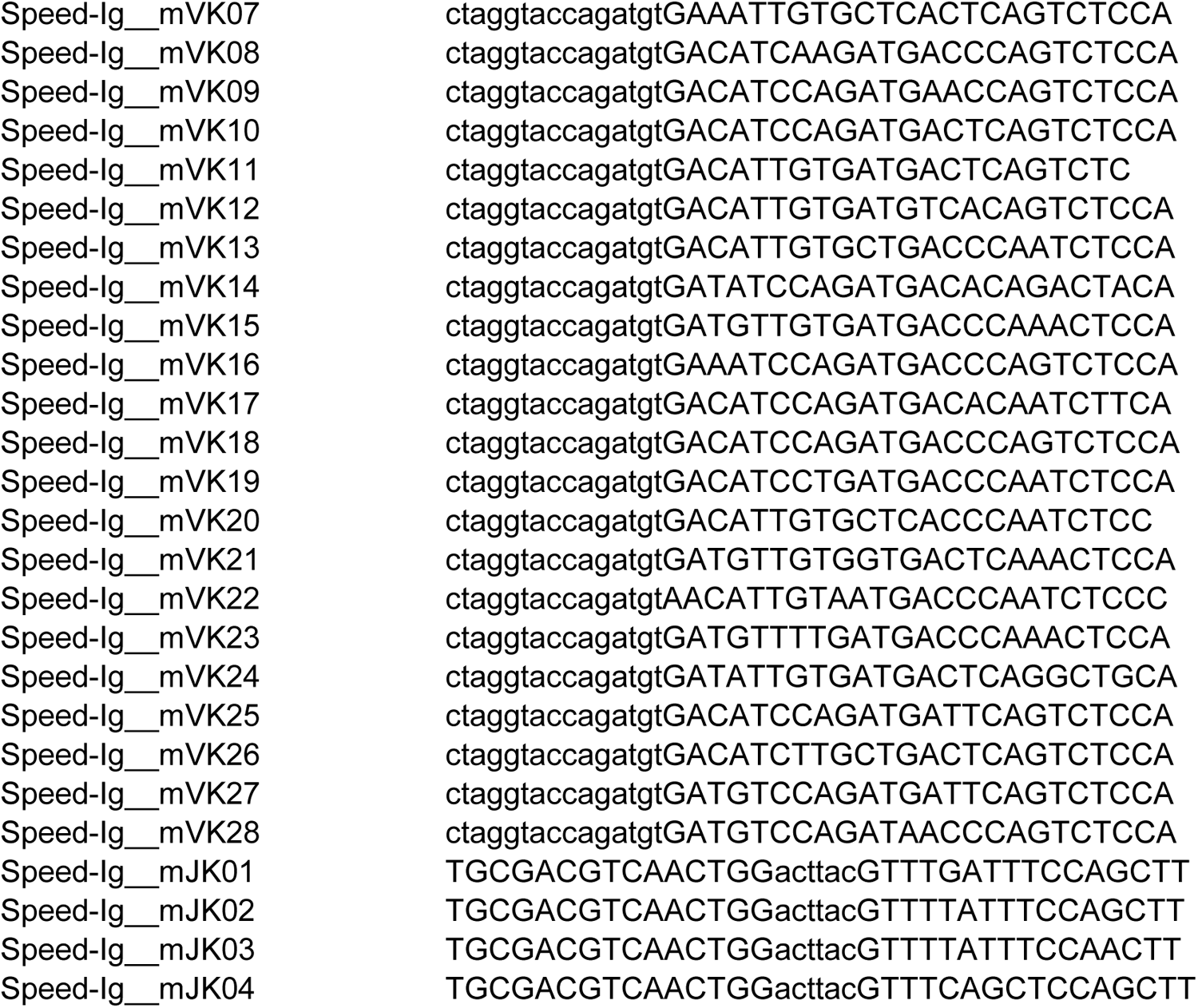
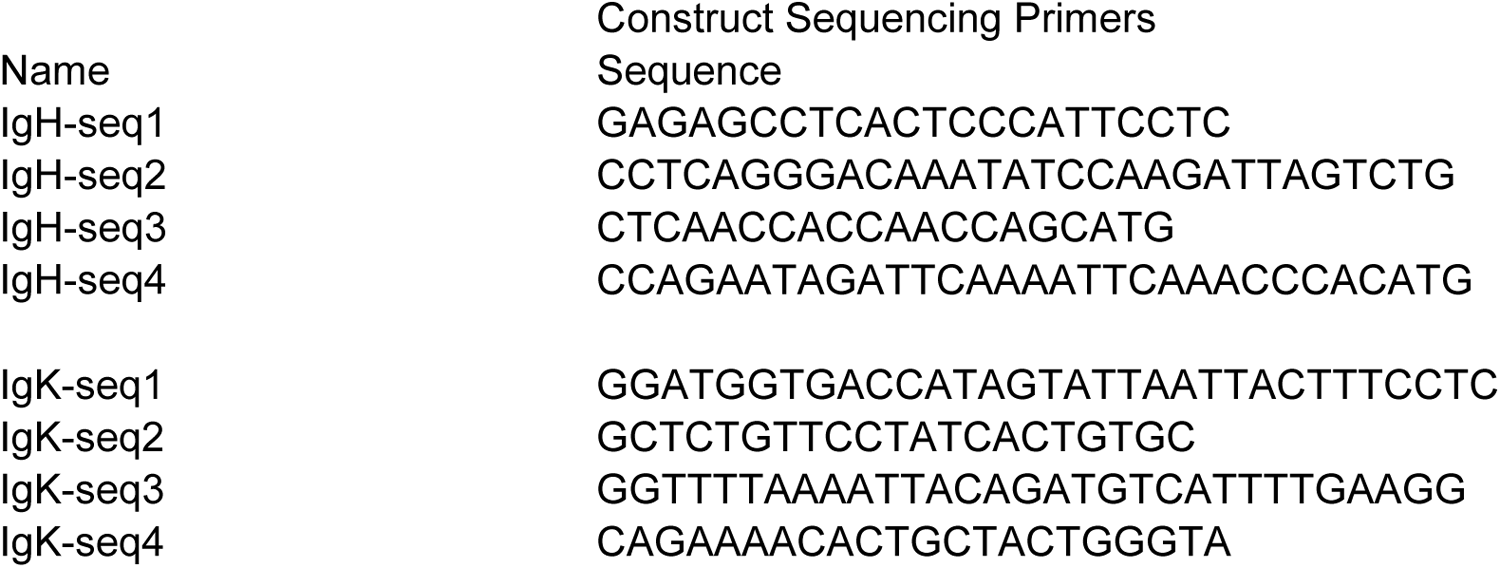
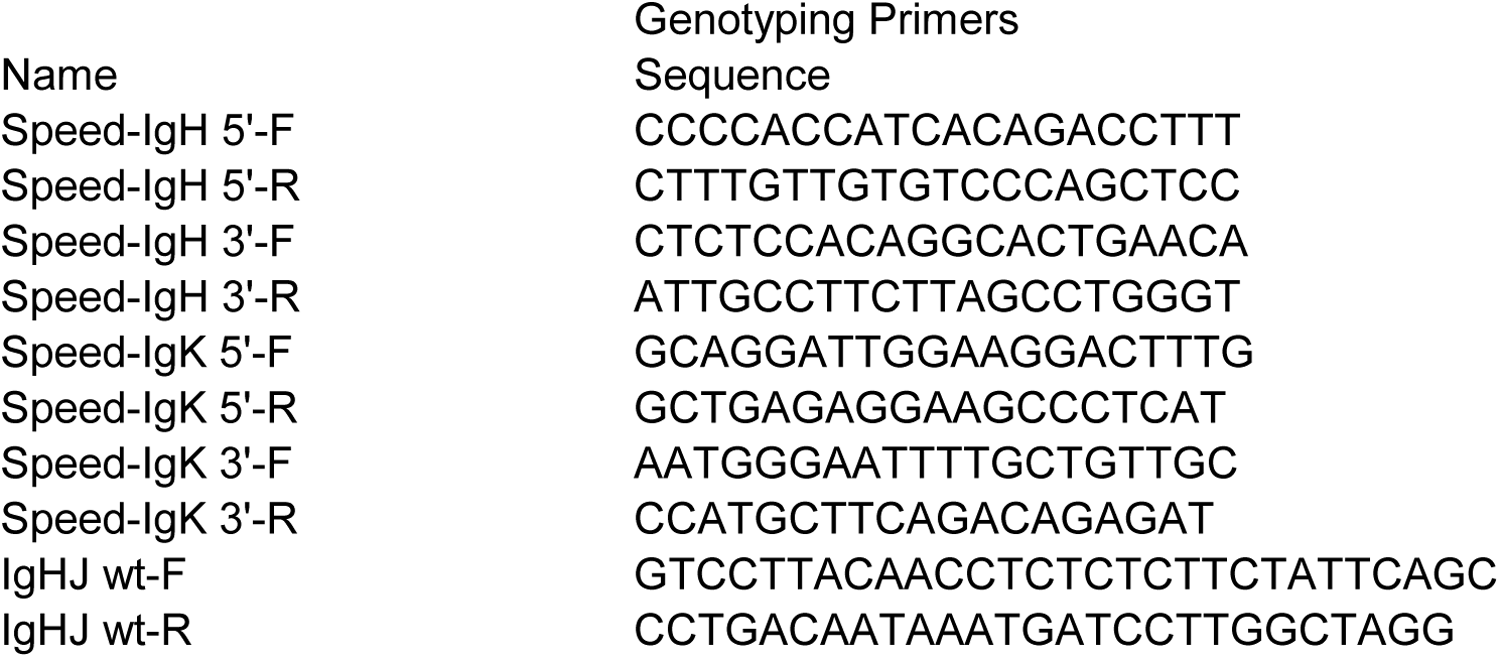
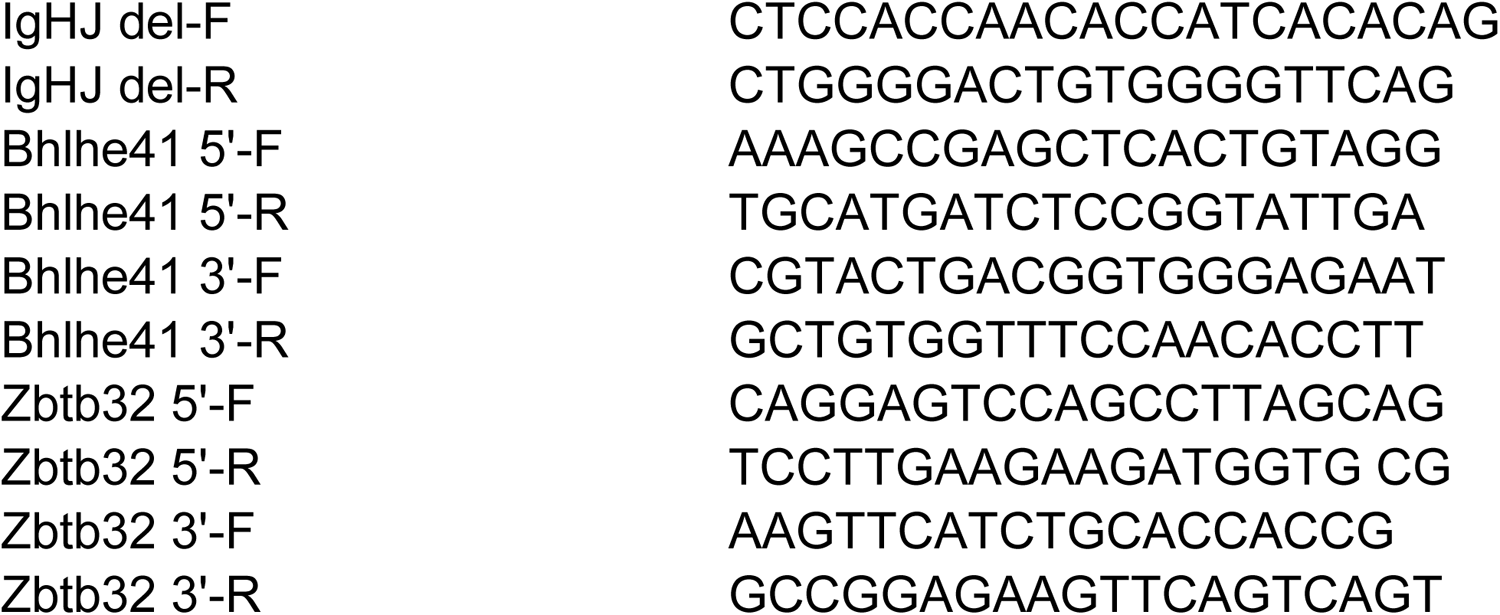
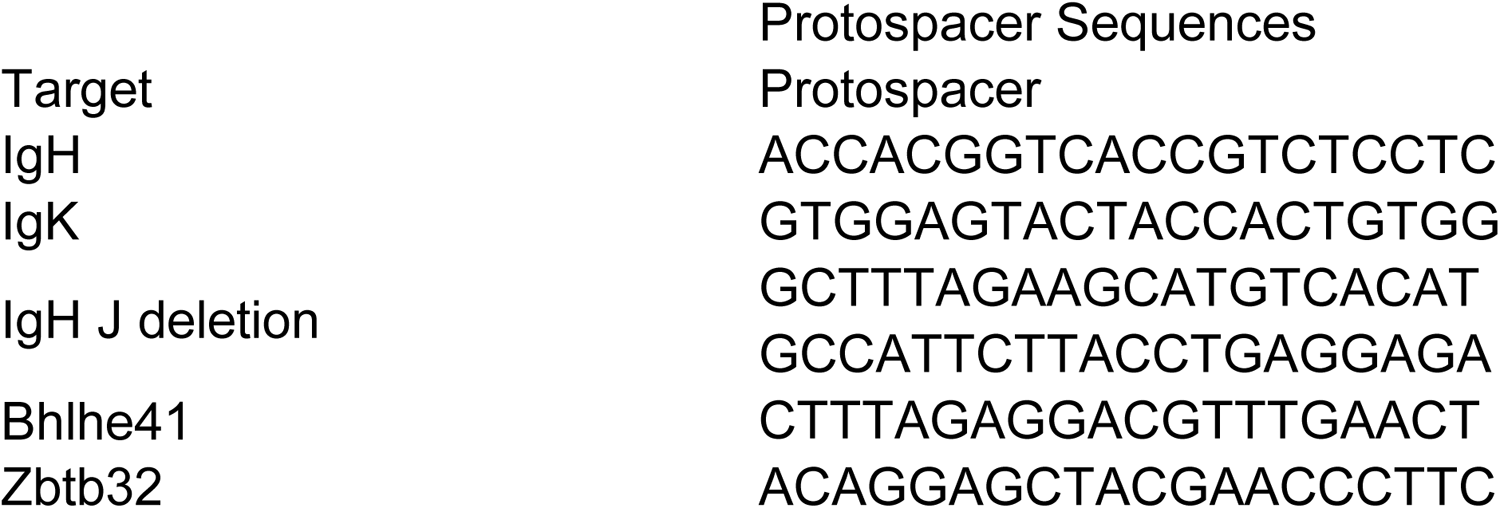
Sequences of primers and crRNAs.

